# Diverse phenotypic responses and phosphate content in foxtail millet genotypes under greenhouse and field conditions

**DOI:** 10.1101/607515

**Authors:** S. Antony Ceasar, M. Ramakrishnan, K. K. Vinod, G. Victor Roch, Hari D. Upadhyaya, Alison Baker, S. Ignacimuthu

## Abstract

Phosphorous (P) is an important macronutrient for the growth of all agricultural crops. This study reports phenotype analysis for P responses in field (two different seasons, monsoon and summer) and greenhouse, using 54 genotypes of foxtail millet (*Setaria italica*) under P-fertilized (P+) and unfertilized (P-) conditions. Variation was seen for plant height, leaf number and length, tillering ability and seed yield traits. Genotypes ISe 1234 and ISe 1541 were P+ responders, and the genotypes ISe 1181, ISe 1655, ISe 783 and ISe 1892 tend more towards low P tolerance for total seed yield. Genotypes that performed well under P-conditions were almost as productive as genotypes that performed well under P+ conditions suggesting some genotypes are well adapted to nutrient-poor soils. In the greenhouse, significant variation was seen for root hair density and root hair number and for fresh and dry weights of shoot and root under P-stress. However, there was not much difference in the shoot and root total P and inorganic phosphate (Pi) levels of five selected high and low responding genotypes. In the root and leaf tissues, total P and Pi contents of five high responding genotypes were higher than the five low responding genotypes.

**Highlight:** Enormous phenotypic and phosphate content variation of foxtail millet under low-phosphate supply in greenhouse and natural field conditions identifies genotypic plasticity for future breeding for improved P use efficiency.

## Introduction

Phosphorous (P) plays essential functions in living systems and it is an irreplaceable element for crop production. Plants uptake P from the soil as inorganic phosphate (Pi) and the process is affected by external factors such as soil pH, microbial activity, cationic abundance and presence of mycorrhizal fungi (*Schachtman et al*., 1998). Although soils are rich in P, paradoxically it remains highly limiting for plant growth due to chemical fixation in organic or in inorganic form. Globally, 5.7 billion hectares of land are affected by Pi-deficiency that significantly hampers crop production (Niu *et al*., 2013). In continuous cropping systems, replacement of depleted nutrients is carried out through external application for fertilizers to sustain crop production. However, projections suggest reserves of rock phosphate, the prime source of inorganic phosphate fertilizers, may only last for the next 200 years (Cordell *et al*., 2009; Sattari *et al*., 2012). The low Pi use efficiency (PUE) of the modern cultivars (around 20%), is a major problem in intensive cropping systems, where the input requirement of phosphatic fertilizers is high, leading to fixation in the soil and/or environmental leaching of the unutilised Pi in water bodies (Vinod & Heuer, 2012). Under circumstances of continuous cropping and Pi fixation, input reduction will exacerbate Pi starvation in crop plants (Vinod, 2015). To tackle this problem, there is a concurrent need for Pi input reduction and crop PUE improvement.

PUE in plants comprises Pi acquisition efficiency (PAE); the extent to which plants acquire Pi from the soil, and Pi utilization efficiency (PUtE); the efficient use of internal P resources. Since both the components are complementary, simultaneous improvement of PAE and PUtE is an important objective for sustainable crop production in order to achieve reduced reliance on external Pi supplementation (Veneklaas *et al*., 2012; Wang *et al*., 2010). Several approaches have been adopted with the goal of improving both the PAE and PUtE of crop plants. These include both transgenic and breeding based approaches (reviewed in Baker *et al*., 2015; Lopez-Arredondo *et al*., 2014). Extensive phenotyping studies have been performed to identify the genotypes with greater PUtE and to map the associated quantitative trait loci (QTL) for use in breeding programs (reviewed in Wiel *et al*., 2015). Most of these studies focus on P starvation tolerance because improvement of P acquisition from soil and utilisation is the prime objective of breeding for PUE in crops. Under P starvation, P homeostasis is sustained through adaptive mechanisms such as improved nutrient foraging in the rhizosphere through root architectural manifestations that include root proliferation and deeper penetration (Lynch & Brown, 2001).

In *Arabidopsis* as well as several crops such as rice, maize, common bean, white lupin, tomato, black mustard and finger millet, root modifications such as primary root length reduction, improved branching, increased lateral root length, enhancement in number of lateral roots and root hair proliferation were reported as Pi starvation responses (Dinkelaker *et al*., 1995; Carswell *et al*., 1996; Borch *et al*., 1999; Kim *et al*., 2008; Lambers *et al*., 2011; Jin *et al*., 2012; Ramakrishnan *et al*., 2017; Niu *et al*., 2013). Additionally, roots play a major role in boosting rhizosphere biology by improving microfloral symbiosis and rhizochemical reactions through root exudates which aids P solubilization and thereby Pi uptake. Root level adaptation augments the innate ability of plants to uptake more Pi from the soil, thereby making it a potential trait targeting improvement in PAE (Lynch & Brown, 2008; Rouached *et al*., 2010; Péret *et al*., 2011). Aerial parts of cereal crops also show key phenotypic modifications in response to low P starvation. These include stunted shoot growth, dark green leaf and reduced yield in oat (Andersson *et al*., 2003), increased primary root length and reduced photosynthesis in rice (He *et al*., 2003; Wissuwa *et al*., 2005), reduction in leaf and primary root growth and photosynthesis in maize (Mollier & Pellerin, 1999) and suppressed shoot growth in sorghum (Yoneyama *et al*., 2007). Reduced shoot growth and lower seed yield were reported in foxtail millet genotype Maxima grown under low P in the greenhouse (Ceasar *et al*., 2014).

Millets are small-seeded cereals in the family Poaceae, mainly cultivated and consumed by people in the tropical world. The millet grains are rich in essential nutrients like calcium, magnesium, zinc and iron and form a major source of nutrition and food security for millions in less developed nations (Ceasar & Ignacimuthu, 2009). Foxtail millet (*Setaria italica*) is widely cultivated in the semi-arid regions of Asia (India, China and Japan) and also in Southern Europe and has the longest history of cultivation among all the millets (Diao & Jia, 2017). Unlike the major cereals, breeding interventions in millets are limited especially using biotechnological tools. Foxtail millet is a genetically amenable model crop since it is a diploid and possesses a relatively small genome of ∼515 Mb. Currently, the genome sequence information of foxtail millet varieties is available (Li & Brutnell, 2011; Bennetzen *et al*., 2012; Zhang *et al*., 2012). Foxtail millet is also closely related to bio-fuel grasses such as switchgrass and Napier grass (Muthamilarasan & Prasad, 2015). Together with its wild relative, green foxtail (*S. viridis*), both *Setaria* species are considered good models for nutrient management in other millets (Huang *et al*., 2016; Ceasar 2018). There are no reports to date exploring the responses to P starvation in foxtail millet, except for a regional report (Kalaghatagi *et al*., 2000). We hypothesise that given similar cultural conditions, with contrasting P levels, those foxtail millet genotypes that fare well under limited P nutrition would be P starvation tolerant and those do well under high P nutrition would be fertilisation responsive. This information would be of value for future breeding programmes. In this study, we report the response of fifty-four genotypes of foxtail millet to P sufficiency (P+) and starvation (P-) under both greenhouse and natural field conditions.

## Materials and methods

### Plant material

Seeds of fifty genotypes used in this study were obtained from the International Crops Research Institute for the Semi-Arid Tropics (ICRISAT), Patancheru, Hyderabad, India. The genotype ‘Maxima1’ (Acc.No: Bs 3875; Welsh Plant Breeding Station, Aberystwyth University, UK) previously used in genetic studies on P response (Ceasar *et al*., 2014; Ceasar *et al*., 2017) and three local genotypes of Southern India (CO5, CO6 and CO7) procured from Tamil Nadu Agricultural University (TNAU), Coimbatore, India) were also included in the trial, and two of them, CO5 and CO7 were used as local checks for comparison. The details of all the fifty-four genotypes are given in **Supplementary Table S1**. These genotypes were evaluated under greenhouse and natural field conditions in two separate experiments conducted in two different seasons. Both the experiments were conducted using a randomized complete block design (RCBD) with two replications.

### Field experimentation

The growth and yield studies were conducted under natural field conditions at Paiyur, Tamil Nadu, India (12°25’ N, 78°13’ E) located at an elevation of 460 meters above sea level and repeated for two years, in the summer season during 2015 (April to July) and in the monsoon season in 2017 (August to November). The experimental station received an average rainfall of ∼105.17 mm in April to July 2015 and ∼188.72 mm in August to November 2017. This location has a history of traditional small millet cultivation for over several hundred years. An unfertilized field that was left barren for several previous seasons was chosen for the study. Prior to the experiment, soil samples were collected from around the field, and the available P was found to be 5.5 mg/kg, which is normally considered as Pi-deficient under natural field conditions (Gemenet *et al*., 2016). Two treatments were maintained, (i) in which no P fertilizer was applied (P-) and (ii) wherein P application was done at double the recommended doses (P+). The crop was raised by sowing the seeds in shallow furrows and plants were grown with a spacing of 30 cm between rows and 10 cm between plants within a row. A block size of 11 x 1 meter was maintained with a spacing of 60 cm between the blocks. Each block consisted of 27 genotypes (25 tests and two checks CO5 and CO7), with one check alternating every three test genotypes. The borders rows were planted with check varieties. There were two such blocks per each P regime (P- and P+). Two replications were maintained. Pi fertilization was done by applying 100g of diammonium phosphate (DAP) only into each of the P+ plots. To compensate the extra N supplied through DAP, 40 g of urea was applied in the P-plots such that the amount of N was the same for both Pi treatments. K was also the same between the Pi treatments. The field was irrigated once a week and the weeding done in every alternate week. The plants were allowed to grow to maturity under normal recommended agronomic practices. Agro-morphological data were collected on per plant basis at maturity on plant height (PH), tiller number (NT), productive tiller number (NPT), leaf number (NL), leaf length (LL), panicle length (LF), cluster number (NC)/panicle, 25 seed weight (25SW), seeds per cluster (SPC) and total seed yield (TSE)/panicle.

### Greenhouse experimentation

Greenhouse experiments were conducted at the Entomology Research Institute (ERI), Loyola College, Chennai, Tamil Nadu, India. Seeds were sown in the 4 L pots (10 seeds/pot) and allowed to germinate and grow in perlite medium (Astrra Chemicals, Chennai, Tamil Nadu, India) supplied with a basal nutrient solution (Ceasar et al., 2014) containing 300 µM of Pi (P+) or 10 µM of Pi (P-). For supplying different Pi regimens in nutrient solutions different ratios of KH_2_PO_4_ and K_2_SO_4_ were used to maintain a constant concentration of potassium. Three replicates were maintained for each of the Pi treatments. The whole experiment was maintained in a greenhouse at 26°C, 16 h light with 85% relative humidity under well-lit and aerated conditions. The shoot dry weight (SDW) and root dry weight (RDW) of 15-day old seedlings for each treatment were observed and combined into a single trait, biomass to avoid reduced variance due to fractionation. Shoot length (SL), root length (RL), root hair density (RHD) and root hair length (RHL) were determined at 28 days. For the analysis of RHD and RHL, the seedlings were carefully removed from perlite, washed twice with distilled water and blotted dry. Root hair measurements were performed according to (Slabaugh *et al*., 2011) with some modifications. About 5 cm from the primary root cap was photographed using a Canon Coolpix digital camera under a Leica Stereo Microscope (Wetzlar, Germany) and measured using ImageJ image processing and analysis software v1.80_112 (Schneider *et al*., 2012).

### Analysis of phosphate content

The total P and Pi contents of the root and shoot tissues of the five most extreme genotypes in each experiment were analysed according to Chiou *et al*., (2006). Selection of extreme genotypes was done independently on both P- and P+ treatments following the same procedure described below.

### Data analyses

Analysis of variance was conducted on the field observations for two seasons using restricted maximum likelihood (REML) approach implemented in ‘lme4’ package in the R statistical environment. The genotypes and season were treated as random factors to estimate the unbiased predictors from the model. Those traits having non-significant genotypic effects were dropped, and the predicted genotype effects of different traits were used for further analyses. To identify the extreme 10% genotypes for the overall phenotypic performance, the trait predictors were ranked in descending order individually for all the traits, so that the highest value gets the rank of one. The cumulative ranking (rank sum) of each of the genotype was computed by adding the ranks of that genotype for all the traits. For instance, under P+, the genotype ISe1387 was ranked 1^st^ for PH, NL, LF, NC and TSE, 2^nd^ for LL, 3^rd^ for SPC and 37^th^ for NPT making its rank sum as 47. Similarly, the rank sums were worked out for the rest of the genotypes. A similar approach was implemented for greenhouse data. From the rank sums, extreme five genotypes were picked having low and high-rank sums, under each category of P- and P+ treatments. Four classes of extreme genotypes were selected, namely, lowest rank sum (high performers) under P+, highest rank sum (low performers) under P+, high and low performers under P-.

### Genotype comparison for P response under field and greenhouse conditions

Field and greenhouse data was used to identify genotypic similarity in P response both under P+ and P-conditions. Before clustering, the data were centred by subtracting the trait means and scaled by dividing by the respective standard deviation. A popular unsupervised centroid-based clustering known as k-means clustering (Hartigan & Wong, 1979) was performed by R using *kmeans* function. The clusters were graphically determined by the package ‘gplots’.

### Cluster analysis based on plasticity

Response to P availability in plants was assumed to affect all the phenotypic traits observed. Therefore, the genotypic P response among different traits (plasticity) was worked out as the percentage difference under both P conditions over the mean P response. Hierarchical clustering was done on the plasticity matrix, using correlations as the distance measure and by performing multiscale bootstrap resampling (Shimodaira, 2004). The *p*-values computed for bootstrap events for each cluster was used for identifying significant clusters. There are two *p*-values computed, BP and AU, BP being the bootstrap probability and AU being the approximately unbiased *p*-value, a better measure of approximation (Shimodaira, 2002, 2004). The computations were performed using the ‘pvclust’ package (Suzuki & Shimodaira, 2006) in R using correlations and average agglomeration with 3000 bootstrap resampling. All the analyses were done using R Studio 1.1.463 running R version 3.5.1.

## Results

### Foxtail millet genotypes show differential response for Pi nutrition under natural field and greenhouse conditions

Analysis of variance revealed that significant genotypic variation existed for all the traits under both P- and P+ conditions except for 25 seed weight, which was dropped from further analyses (**Table 1**). The season and genotype x season component showed non-significant variance. The heritability calculated from the estimated genotypic variance obtained from the vector of genetic effects and the error variance obtained from the vector of residuals showed profound influence of genotypic variance to the total variance for all the traits in the population. Although these estimates are not accurate, they indicate the general trend of influence of different sources of variation to the heritable component of the trait (Gilmour *et al*., 1995).

**Table 1.** Testing for the significance of variance component effects by restricted maximum likelihood (REML) method.

| Traits | Variance |  |  | AIC | Heritability |
| --- | --- | --- | --- | --- | --- |
|  | Genotype | Season | Genotype x Season |  |  |
| H-PH | 269.5* | 7.50 | 4.05 | 2701.53 | 0.95 |
| H-NPT | 5.68* | 0.11 | 0.54 | 1638.07 | 0.91 |
| H-NL | 1.21* | 0.05 | 0.00 | 1251.44 | 0.87 |
| H-LL | 37.8* | 0.34 | 0.64 | 2085.77 | 0.95 |
| H-LF | 16.23* | 0.48 | 1.05 | 1702.24 | 0.97 |
| H-NC | 270.73* | 4.21 | 5.98 | 2446.55 | 0.97 |
| H-SPC | 379.2* | 4.52 | 34.4 | 2310.76 | 0.95 |
| H-TSE | 41.12* | 0.32 | 0.18 | 880.99 | 0.99 |
| L-PH | 511.32* | 5.46 | 2.05 | 2307.75 | 0.98 |
| L-NPT | 6.54* | 0.32 | 0.02 | 1503.11 | 0.92 |
| L-NL | 1.99* | 0.03 | 0.00 | 1120.4 | 0.93 |
| L-LL | 52.24* | 0.26 | 0.04 | 1918.04 | 0.95 |
| L-LF | 15.6* | 0.34 | 0.47 | 1531.03 | 0.96 |
| L-NC | 344.5* | 14.04 | 9.74 | 2104.48 | 0.97 |
| L-SPC | 285.97* | 1.62 | 6.61 | 1905.93 | 0.98 |
| L-TSE | 45.76* | 0.42 | 0.21 | 656.35 | 0.99 |

\*Significant at $p < 0.05$ by chi-square test. AIC, Akaike information criterion. Trait names are prefixed H- for P+ response and L- for low P response. PH, plant height in cm; NT, number of tillers; NPT, number of productive tillers; NL, number of leaves; LL, length of leaf in cm; LF, length of panicle in cm; NC, number of clusters/panicle; SPC, seeds per cluster; TSE total seed yield in g.

Based on the cumulative ranking, genotypes ISe 1387 and ISe 1687 were in the top 5 in both Pi fertilized and unfertilised field plots. Other genotypes that performed well under both P- and P+ conditions were CO7 (4^th^, 7^th^), ISe132 (10^th^, 2^nd^), ISe1851 (7^th^, 3^rd^) and ISe 869 (5^th^, 14^th^). Other top genotypes were ISe 663 (3^rd^ under P-) and ISe 132 and ISe 907 (2^nd^ and 4^th^ under P+), **(Figure 1A)**. Maxima1 was the poorest among all the genotypes under P-, followed by ISe1335, ISe1234, ISe1037 and ISe1302. Maxima1, ISe1335, and ISe1234 were also in the bottom 5 under P+ treatment.

**Figure 1.**
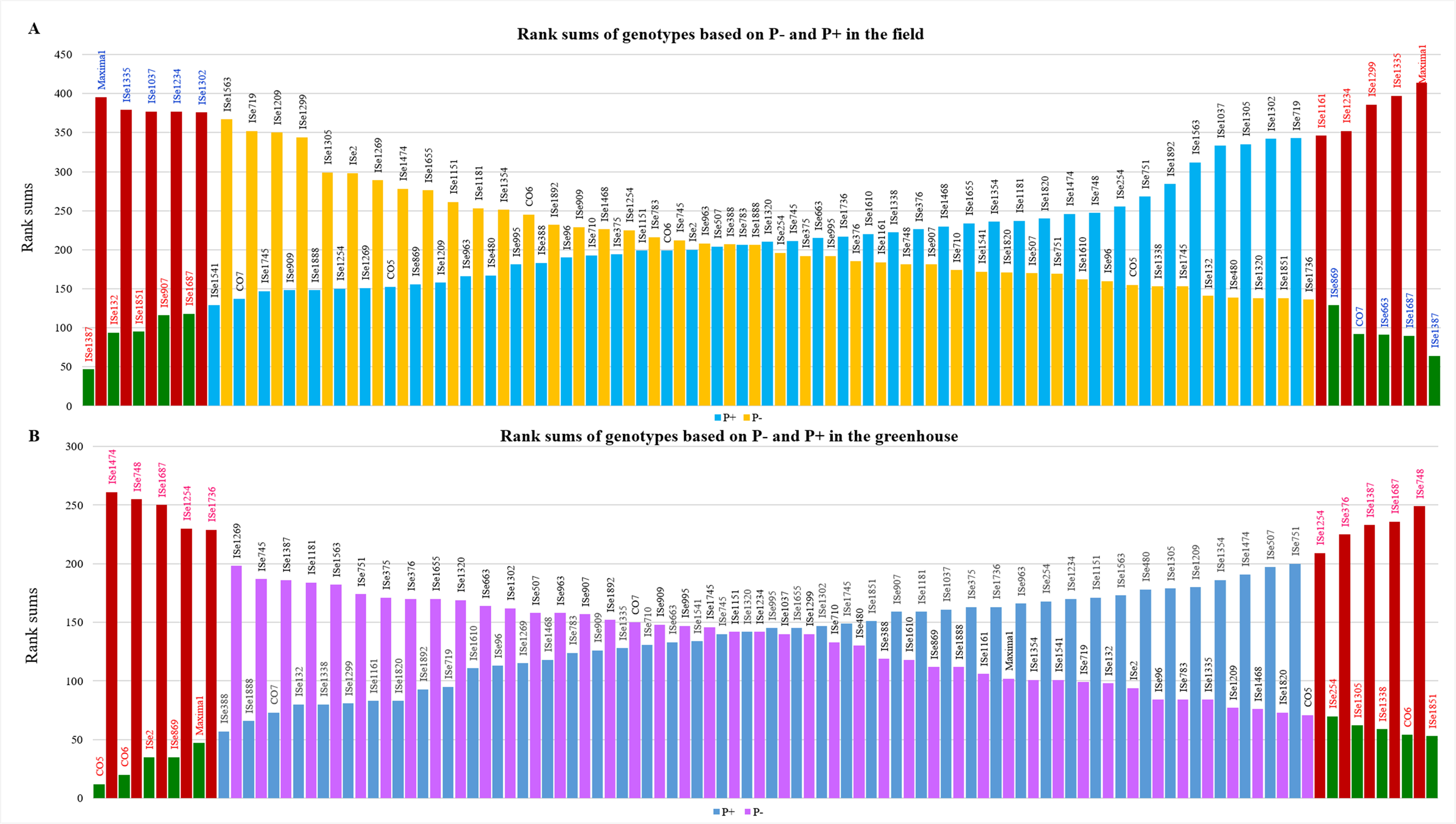
Rank sums of genotypes based on (A) field and (B) greenhouse evaluation. The 10 extreme genotypes are selected under each P regime, P+ and P-. P+ ranks are ordered from poor (having high-rank sum) to good (having low-rank sum). P ranks are ordered from good (low-rank sum) to poor (high-rank sum). Five genotypes with the lowest rank sum in each group are selected as best genotypes (green), and five genotypes each with highest rank sum are selected as poor genotypes (red).

The means for the field experimentation clearly illustrated the response of genotypes for each trait analysed **(Table 2 and Supplementary Table S2)**. Statistically significant differences were seen for all the traits when the good and poor performing genotypes based on the rank sums for the P+ plot were compared. In the case of P-conditions also, conspicuous differences were recorded for all the traits. However, when the selection was done for extreme five genotypes in both the directions in both the treatments, differences in phenotypic performance was found reduced. The inflorescence images depicting the variation in some of the genotypes with varied responses to P fertilization is given in **Figure 2**.

**Table 2.**
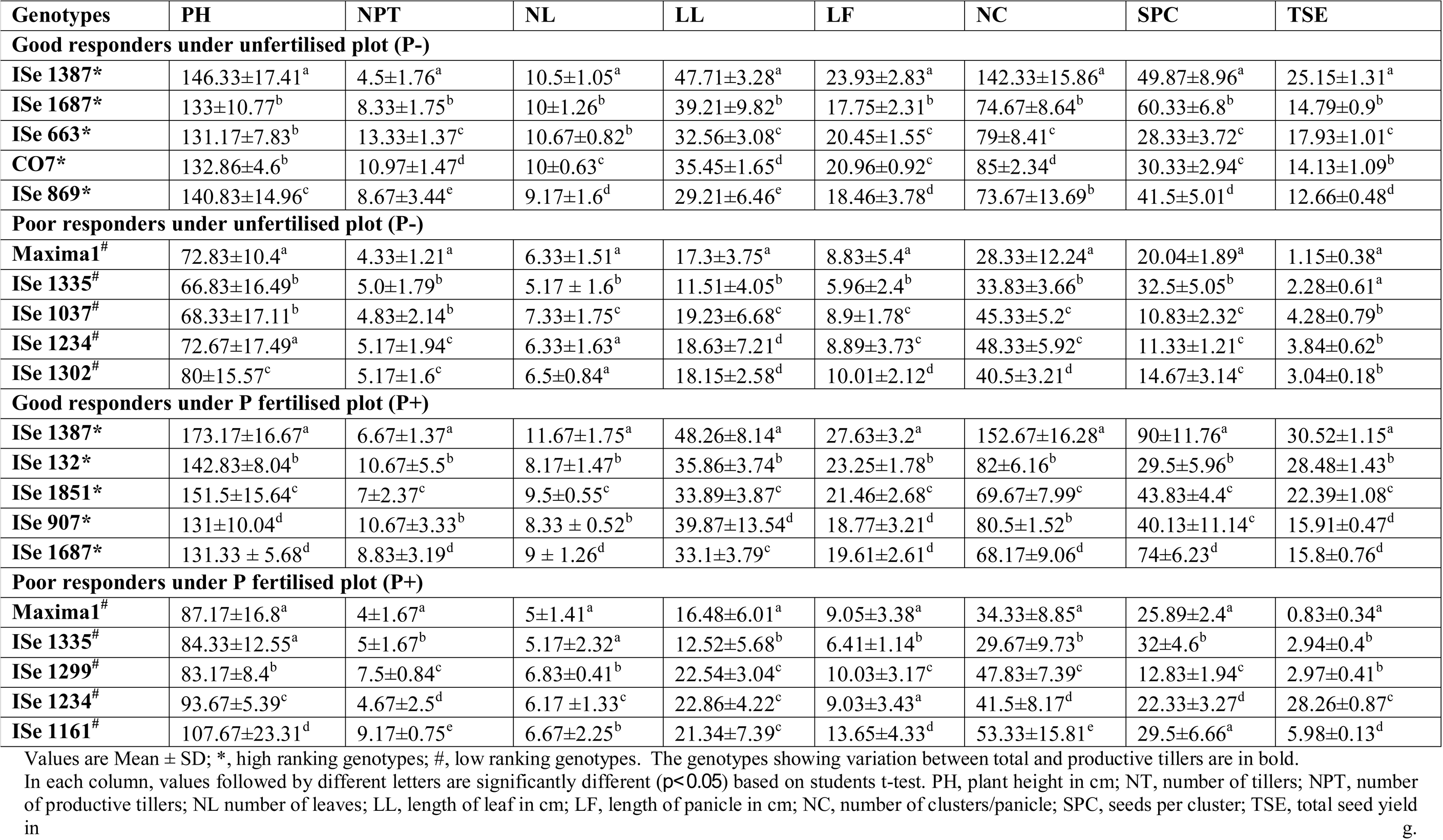
Mean values of the five high and low responding genotypes of foxtail millet in growth assays under unfertilised (P-) and P fertilised (P+) natural field conditions

**Figure 2.**
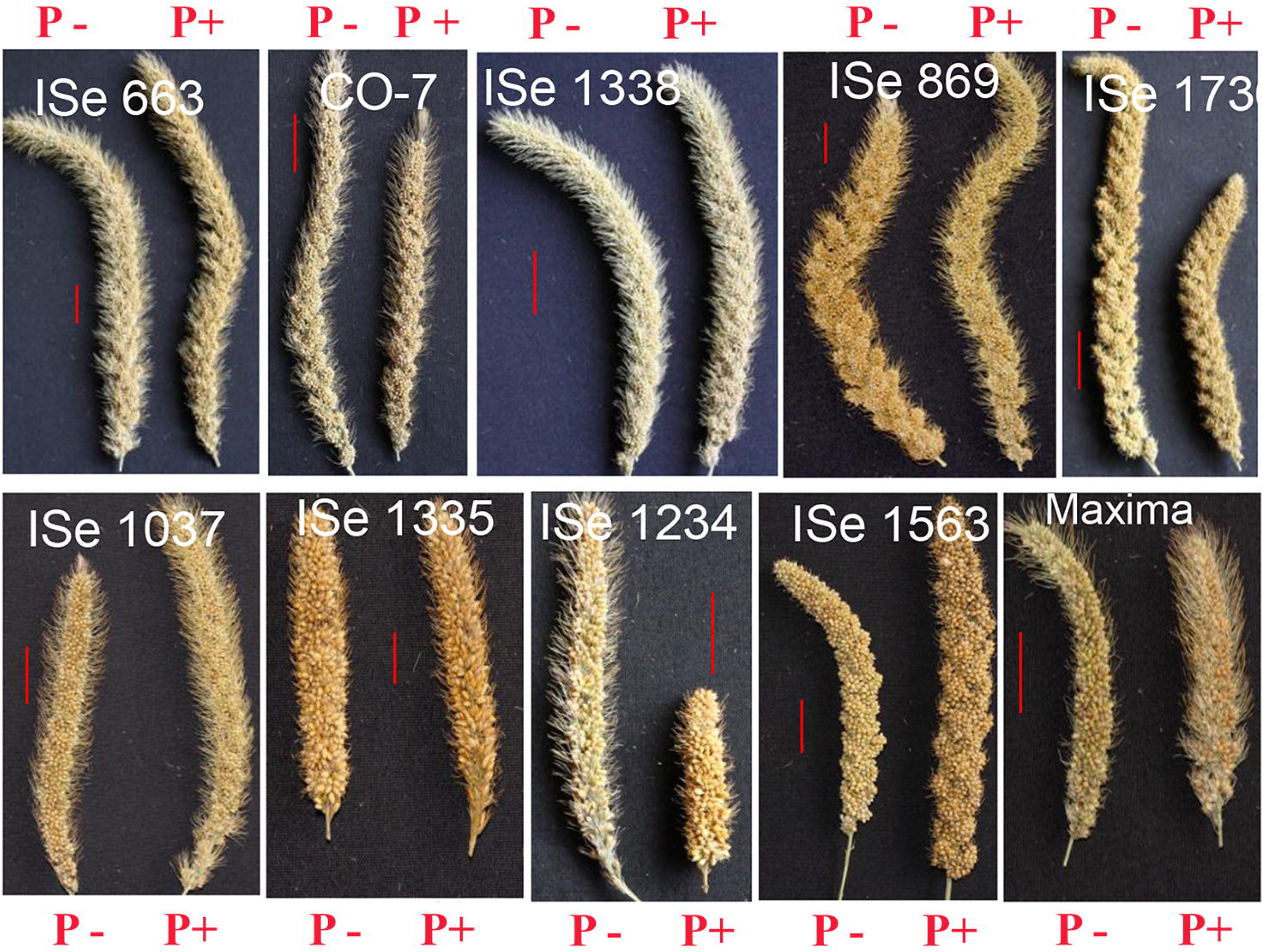
Flowers with mature seeds of selected genotypes (good, intermediates and poor responding) of foxtail millet grown in a natural field Pi fertilized (P+) and unfertilized (P-) soil conditions (P-, low phosphate; P+, high phosphate). The genotypes in the top row are ISe 663, CO7, ISe 1338, ISe 869 and ISe 1736 and bottom row are ISe 1037, ISe 1335, ISe 1234, ISe 1563 and Maxima1.

Greenhouse grown plants also showed a high degree of variation with respect to growth on differing levels of Pi (**Supplementary Table S3)**. Genotype ISe 1851 had the lowest rank sum (high performer) under P-, this was followed by CO6, ISe 1338, ISe 1305 and ISe 1254. (**Figure 1B**). Genotypes ISe 1474 and ISe 748 had the highest (low performer) rank sum under P-. Under high phosphate, the cumulative rank order was very different, though CO6 was 2nd. Genotype ISe 748 performed poorly under P+ as well as P- (**Figure 1B**). Collectively, the high performing genotypes under P-conditions had significantly higher biomass, SL, RL, RHD and RHL than low performers (**Table 3**.) The low performer ISe 1687, registered significantly higher RHD and RHL than other low performers under P-condition. ISe 1687 was interesting because despite RHL and RHD that were comparable to the high performers all other responses were poor, probably because the roots themselves were very small. Under P+ conditions, the five low performers were comparable to the five low performers under P-conditions for all parameters. Genotypes ISe748, ISe1254 and ISe1687 were common to both treatments as low performers. ISe748, ISe1254 and ISe1736 grown under P+ treatment as well as under P-condition lacked root hairs (**Table 3** and **Figure 3).** In contrast, the root hairs in the other genotypes (ISe 1851, ISe 1305, ISe 1209 and ISe 1387) shown in **Figure 3** were sparse or undiscernible under P+ conditions but were increased in length and number in P-conditions. The high responding genotypes under P+ were different from the high responding genotypes under P-with the exception of the local genotype CO6 which was ranked 2^nd^ under both P- and P+ (**Figure 1B**). It attained 90 % of the SL and 76 % of the biomass under the P-conditions as compared to P+. For the root traits under P-, CO6 attains 69 % of the RL, 209 % of the RHD and 113 % of the RHL when compared to P+ conditions.

**Table 3.** Mean values for the five high and low responding genotypes of foxtail millet in growth assays under low (P-) and high (P+) in greenhouse conditions

| Genotypes | BIO | SL | RL | RHD | RHL |
| --- | --- | --- | --- | --- | --- |
| **Good responders under P+** |  |  |  |  |  |
| **CO-5\*** | 8.00±1.00<sup>a</sup> | 21.73±7.98<sup>a</sup> | 13.13±0.87<sup>a</sup> | 41.67±4.04<sup>a</sup> | 3.16±0.01<sup>a</sup> |
| **CO-6\*** | 10.47±0.15<sup>b</sup> | 15.87±5.24<sup>b</sup> | 19.60±0.82<sup>b</sup> | 21.67±3.06<sup>b</sup> | 3.26±0.00<sup>a</sup> |
| **ISe 2\*** | 7.10±1.00<sup>c</sup> | 17.30±0.61<sup>c</sup> | 15.00±1.00<sup>c</sup> | 13.67±1.53<sup>c</sup> | 3.04±0.04<sup>a</sup> |
| **ISe 869\*** | 6.70±0.62<sup>c</sup> | 18.07±5.97<sup>c</sup> | 8.80±1.25<sup>d</sup> | 23.33±2.52<sup>d</sup> | 3.56±0.07<sup>b</sup> |
| **Maxima1\*** | 15.30±1.00<sup>d</sup> | 10.20±2.35<sup>d</sup> | 10.60±3.08<sup>e</sup> | 24.67±3.51<sup>d</sup> | 2.57±0.04<sup>c</sup> |
| **Poor responders under P+** |  |  |  |  |  |
| **ISe 748#** | 2.13±0.60<sup>a</sup> | 2.60±0.92<sup>a</sup> | 5.17±0.40<sup>a</sup> | 0.00±0.00 | 0.00±0.00 |
| **ISe 1687#** | 2.80±2.27<sup>b</sup> | 5.17±1.67<sup>b</sup> | 5.40±0.26<sup>ab</sup> | 1.00±1.73<sup>a</sup> | 0.49±0.00<sup>a</sup> |
| **ISe 1387#** | 3.00±0.56<sup>b</sup> | 4.53±0.95<sup>b</sup> | 6.70±0.62<sup>c</sup> | 0.00±0.00 | 0.00±0.00 |
| **ISe 376#** | 3.50±0.30<sup>c</sup> | 8.43±2.40<sup>c</sup> | 5.93±1.55<sup>b</sup> | 0.00±0.00 | 0.00±0.00 |
| **ISe 1254#** | 6.10±1.15<sup>d</sup> | 2.93±0.51<sup>a</sup> | 4.87±0.67<sup>ab</sup> | 1.67±2.89<sup>a</sup> | 0.40±0.00<sup>a</sup> |
| **Good responders under P-** |  |  |  |  |  |
| **ISe 1851\*** | 7.03±3.67<sup>a</sup> | 9.13±2.81<sup>a</sup> | 12.43±3.77<sup>a</sup> | 51.33±3.21<sup>a</sup> | 4.85±0.10<sup>a</sup> |
| **CO-6\*** | 8.00±0.46<sup>c</sup> | 14.40±3.27<sup>b</sup> | 13.70±1.62<sup>b</sup> | 45.33±3.79<sup>b</sup> | 3.70±0.11<sup>b</sup> |
| **ISe 1338\*** | 6.66±0.32<sup>a</sup> | 15.90±2.25<sup>c</sup> | 16.10±2.21<sup>c</sup> | 49.33±1.53<sup>a</sup> | 3.75±0.12<sup>b</sup> |
| **ISe 1305\*** | 6.07±2.15<sup>b</sup> | 13.00±0.50<sup>d</sup> | 9.50±4.74<sup>d</sup> | 50.67±4.51<sup>a</sup> | 6.24±0.05<sup>c</sup> |
| **ISe 254\*** | 5.87±0.81<sup>b</sup> | 10.43±4.56<sup>e</sup> | 9.13±1.21<sup>d</sup> | 44.00±3.61<sup>b</sup> | 8.01±0.10<sup>d</sup> |
| **Poor responders under P-** |  |  |  |  |  |
| **ISe 1474#** | 2.07±0.59<sup>a</sup> | 4.50±2.18<sup>b</sup> | 2.47±0.45<sup>a</sup> | 1.20±0.00<sup>a</sup> | 1.06±0.18<sup>a</sup> |
| **ISe 748#** | 2.90±0.36<sup>b</sup> | 5.90±1.91<sup>b</sup> | 1.67±0.76<sup>b</sup> | 0.00±0.00 | 0.00±0.00 |
| **ISe 1687#** | 2.40±1.01<sup>a</sup> | 5.17±2.02<sup>b</sup> | 3.50±1.37<sup>c</sup> | 31.00±3.00<sup>b</sup> | 2.37±0.03<sup>b</sup> |
| **ISe 1254#** | 4.43±0.78<sup>c</sup> | 6.67±0.38<sup>c</sup> | 4.73±0.25<sup>d</sup> | 0.00±0.00 | 0.00±0.00 |
| **ISe 1736#** | 3.90±0.96<sup>d</sup> | 5.40±1.89<sup>b</sup> | 7.10±1.21<sup>e</sup> | 0.00±0.00 | 0.00±0.00 |

Values are Mean ± SD; \*= High ranking genotypes; # = Poor ranking genotypes. Within each column, values followed by different letters are significantly different ( $p<0.05$ ) based on students t-test.
BIO, seedling biomass in mg; SL, shoot length in cm; RL, root length in cm; RHD, root hair density per 10µm length, RHL, root hair length in µm.

**Figure 3.**
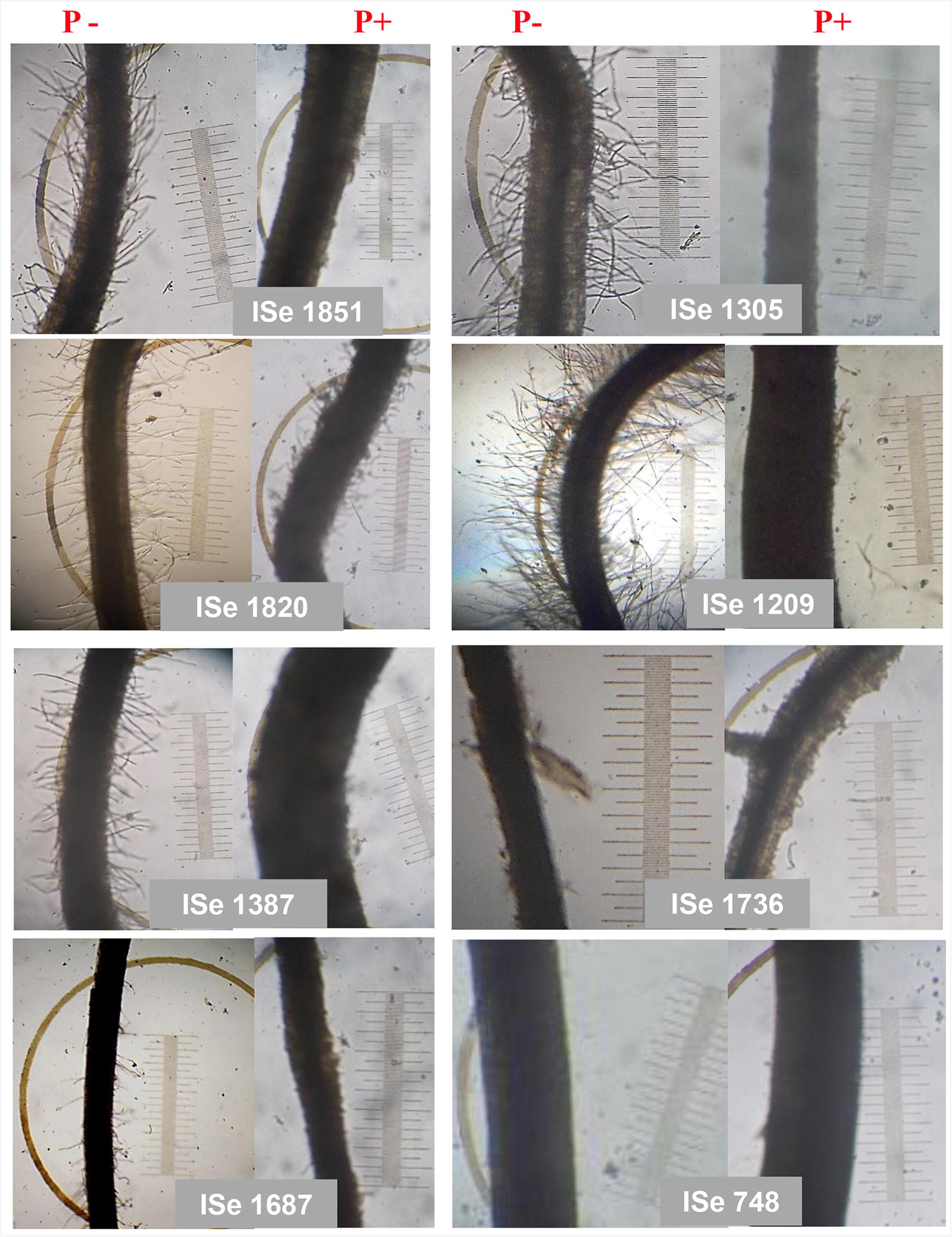
Root hair images of selected genotypes of foxtail millet showing a response to P- and P+ in the greenhouse. The image was taken after 15 days of growth under P- and P+ in the greenhouse. The genotypes ISe 1851 and ISe 1305 are the top in the cumulative ranking and high responding genotypes for root hair formation under P-condition. The genotypes ISe 748, ISe 1687 and ISe 1736 are low performers under P-. The genotype ISe 1387 is a low performer under P+. The genotype ISe 1820 is an intermediate responder in both P- and P+.

ISe 1387 which was the highest ranking genotype under field conditions in both Pi regimes, ranked low when grown under greenhouse conditions (**Figure 1A, B**). Maxima1, which showed a poor field performance under both fertilised and unfertilised conditions, was the fourth best performer under P+ treatments, and mid-ranking under P- in the greenhouse screen. Similarly, CO6 which performed very well under both P+ and P- in the greenhouse experiments indicated a moderate response under field conditions (37^th^ under P-conditions and 24^th^ under P+ conditions). ISe 1851 performed well under both field as well as greenhouse assays and was the most consistent high-responding genotypes in the study. Other high performers were CO7 and ISe 869. Further, the genotype CO5 which was placed at the 13^th^ position in both P- and P+ soil, was placed at the first position under P+ and 6^th^ position under P-situations under greenhouse evaluation.

### P contents of high-responding genotypes were higher than low performers under field and greenhouse conditions

The total P and Pi contents were assayed in the leaf and root tissues of each of the five high and low responding genotypes under P- and P+ conditions, grown in the greenhouse (**Figure 4**) as well as in the field (**Figure 5**). Good responders under the P+ condition in the glasshouse generally had much higher (1.5-3 fold higher) total leaf P and leaf Pi contents than low performers. Root total P and Pi was also higher although there was more variation and some overlap with some of the poor responders such as ISe748 and ISe 1387 (**Figure 4A and B**). The low responding genotype ISe 748 had the same level of Pi in root tissues as some of the high-responding genotypes (CO5 and ISe 869) but much less in the shoot tissues. This may suggest that ISe 748 had low ability in exporting Pi from the root to shoot tissues (compare **Figure 4A and B**). Under P-, the high responding genotypes maintained higher levels of both root and shoot P compared to poor responders (**Figure 4C and D**). As expected, in both high and low performers, more plant Pi is present under P+ conditions than under P-conditions (compare **Figure 4A and C and 4B and D**), but the good responders on P-maintained similar levels of tissue P to poor responders on P+ suggesting higher PAE (compare **Figure 4B and C**).

**Figure 4.**
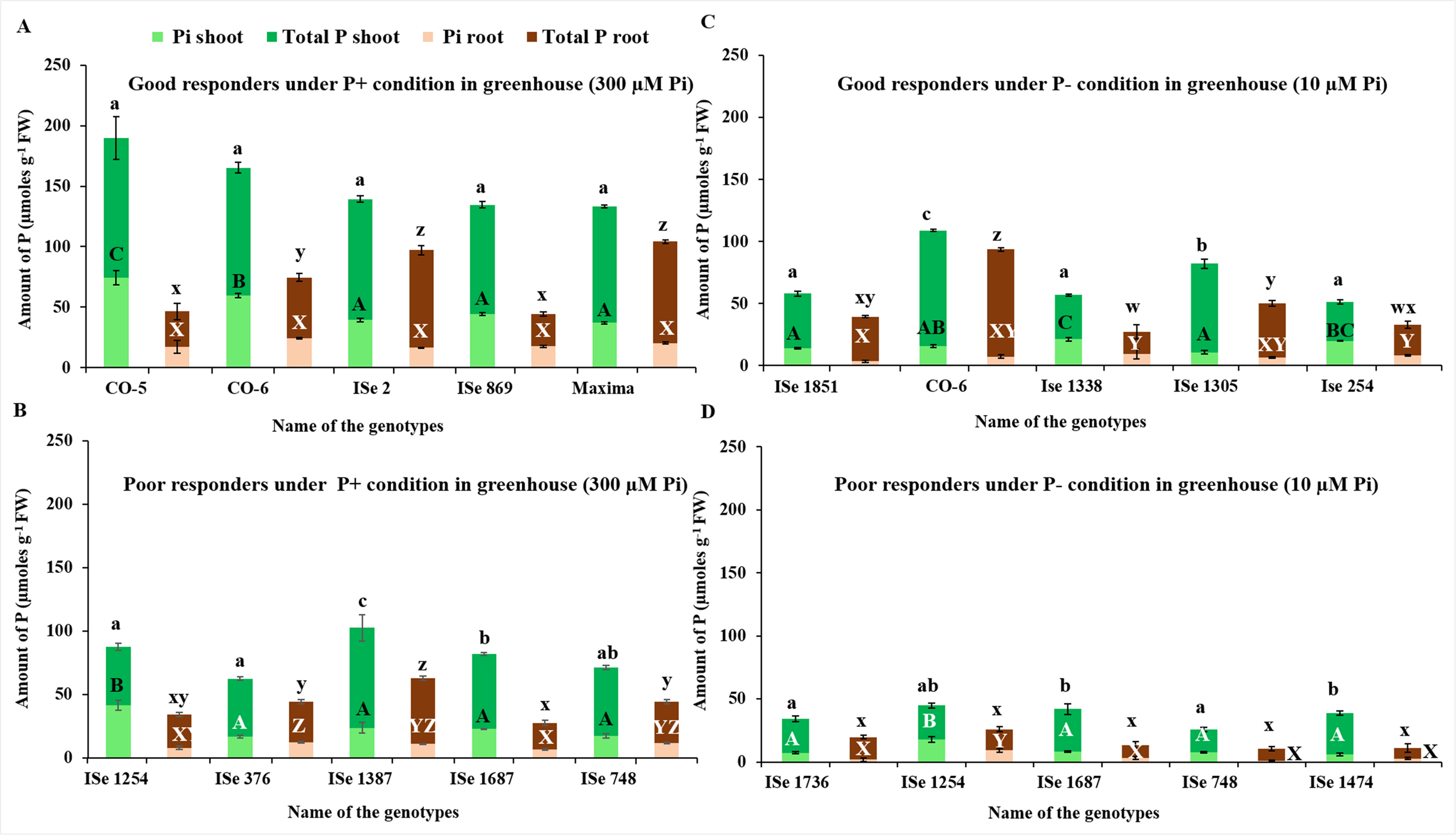
Inorganic (green) and total P assay (brown) for the five each of high performers and low performers of foxtail millet genotypes under P- (10 µM Pi) and P+ (300 µM Pi) conditions in greenhouse conditions. A, high performers under P+; B, low performers under P+; C, high performers under P-; D, low performers under P-. The letters over the bars indicate statistically significant difference if the letters are different as p<0.05. The vertical lines indicate error bars.

**Figure 5.**
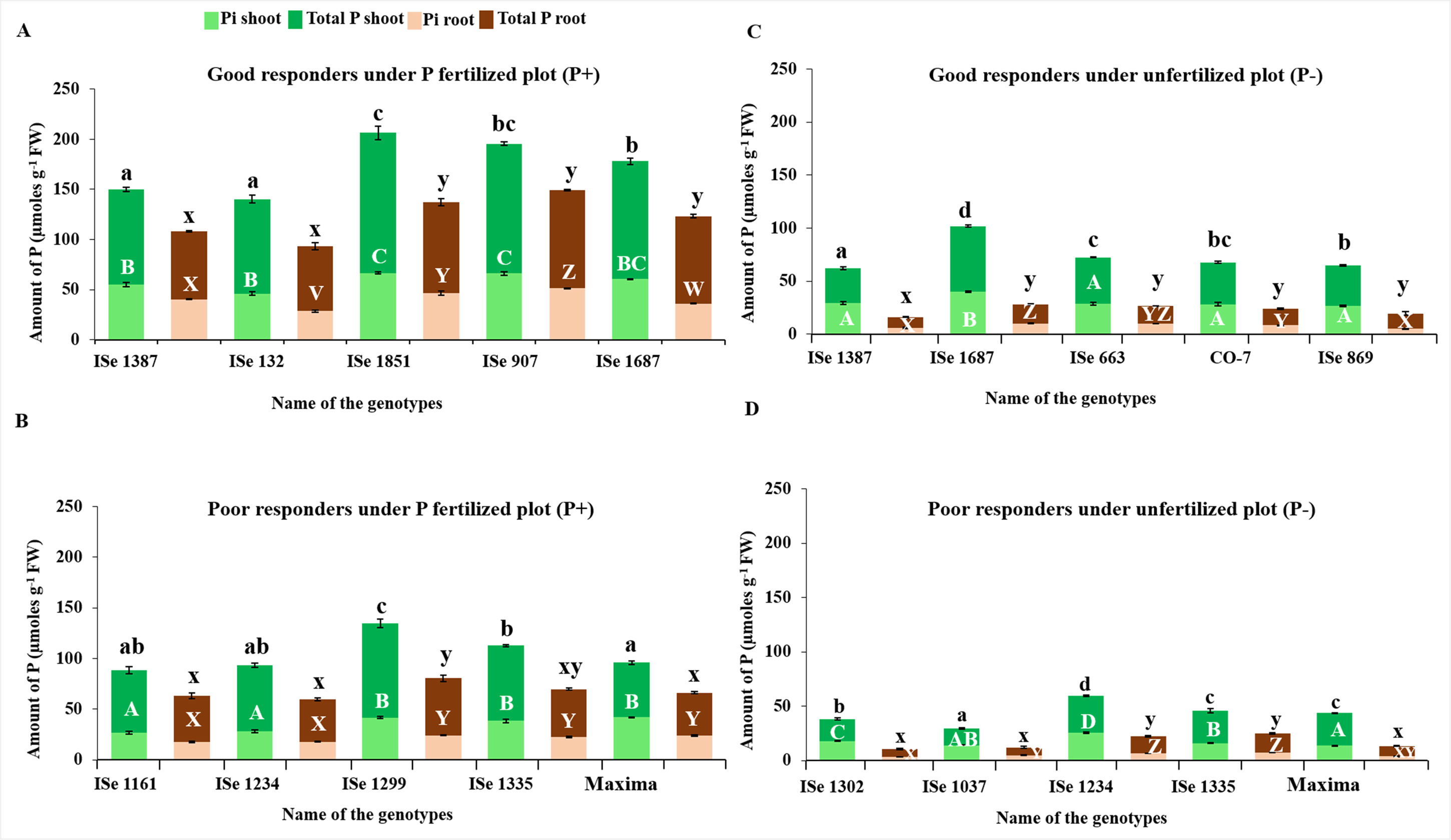
Inorganic (green) and total P assay (brown) for the five each of high performers and low performers of foxtail millet genotypes under unfertilized (P-) and P fertilized (P+) conditions in the natural field. A, high performers under P+; B, low performers under P+; C, high performers under P-; D, low performers under P-. The letters over the bars indicate statistically significant difference if the letters are different as p<0.05. The vertical lines indicate error bars.

Under field conditions, the best performers had higher (>50 %) total and Pi content than the poor performers (**Figure 5A and B**). The levels of shoot total P and Pi were similar between the good responders under glasshouse and field conditions but root total P and Pi was higher in the best performers under field conditions compared to glasshouse (**Figure 4A and Figure 5A**). Most of the good responding genotypes grown under P-condition in the field maintained similar levels of total P and Pi in shoots compared to poor responders (compare **Figure 5C and 5D**). The good responders on P-maintained similar levels of leaf Pi to poor responders on P+ but root total P and Pi was much lower, again pointing to effective export from root to shoot (**Figure 5B and C**). For example, the high responding genotype ISe 1687 had more than 100 µmoles/gm total P which was significantly greater than other high performers and all low performers under P-treatment.

Interrelations between P content among the extreme genotypes showed that shoot and root Pi contents were significantly correlated to total P in the respective tissues both under field and greenhouse screening **(Table 4)**. Under greenhouse situations, Pi content of shoot tissues was also related to root Pi content but not to total root P content. Overall, under field situations, both the root Pi and shoot Pi were correlated as well as to the total P content in both the plant tissues. Conspicuously, there was good agreement between P content of the same genotypes grown under field and greenhouse conditions in both the root and shoot tissues. Analysis of the correlations between tissue-specific P content assayed under field and greenhouse, within different categories of genotypes falling within the four response classes indicated discernible deviation from the overall pattern. Shoot Pi of the good and poor responders under P-condition showed no apparent relation between field and greenhouse conditions, whereas the same under P+ conditions showed significant but negative relations. Negative correlations were also recorded between field and greenhouse-based assays of total shoot P as well as root Pi, under P+ regime. In poor performers, however, total root and shoot P showed a significant positive correlation between field and greenhouse data.

**Table 4.** Interrelations between P content in shoot and roots among the extreme genotypes showing P response under greenhouse (upper diagonal) and field (lower diagonal) conditions. The diagonal values (in bold) are correlations between screen house and field parameters. Values shown against genotype response groups are the correlations between field and greenhouse parameters under each category.

|  | *Pi shoot* | *Total P shoot* | *Pi root* | *Total P root* |
| --- | --- | --- | --- | --- |
| Pi shoot | **0.705\*** | 0.761* | 0.820* | 0.294 |
| Total P shoot | 0.949* | **0.842\*** | 0.790* | 0.730* |
| Pi root | 0.940* | 0.953* | **0.808\*** | 0.526* |
| Total P root | 0.919* | 0.971* | 0.978* | **0.501\*** |
| Good P- | -0.157 | 0.747* | 0.407 | 0.418 |
| Poor P- | -0.250 | 0.215 | -0.190 | -0.692* |
| Good P+ | -0.668* | -0.746* | -0.878* | 0.237 |
| Poor P+ | -0.507* | 0.870* | 0.066 | 0.698* |

\*significant as $p < 0.05$

### Genotypic plasticity for seed yield identifies low P tolerant genotypes based on P response

Genotypic plasticity, the ability of the plants to withstand the extreme variations in P availability, could identify three sets of genotypes such as low P tolerant, high P tolerant (P+ responders) and intermediates under both P+ and P- conditions. The genotypes ISe 1181, ISe 1655, ISe 783, ISe 1892 were low P tolerant based on seed yield (on the left side of **Figure 6)**. Genotypes ISe 1234, ISe 1541, ISe 1563, ISe 1820, and ISe1888 were P+ responders and hence were identified as high P tolerant types (on the right side of **Figure 6)**. It is of particular interest to understand which genotypes were most responsive to P fertilisation. Although plasticity over the mean performance was low (19.3%), the genotype ISe 1387 showed a particularly strong response to P application in both greenhouse and field conditions, in terms of absolute phenotypic performance (**Figure 4 and 5**). But, this genotype was high P tolerant, not low P tolerant based on plasticity analysis (**Figure 6**).

**Figure 6.**
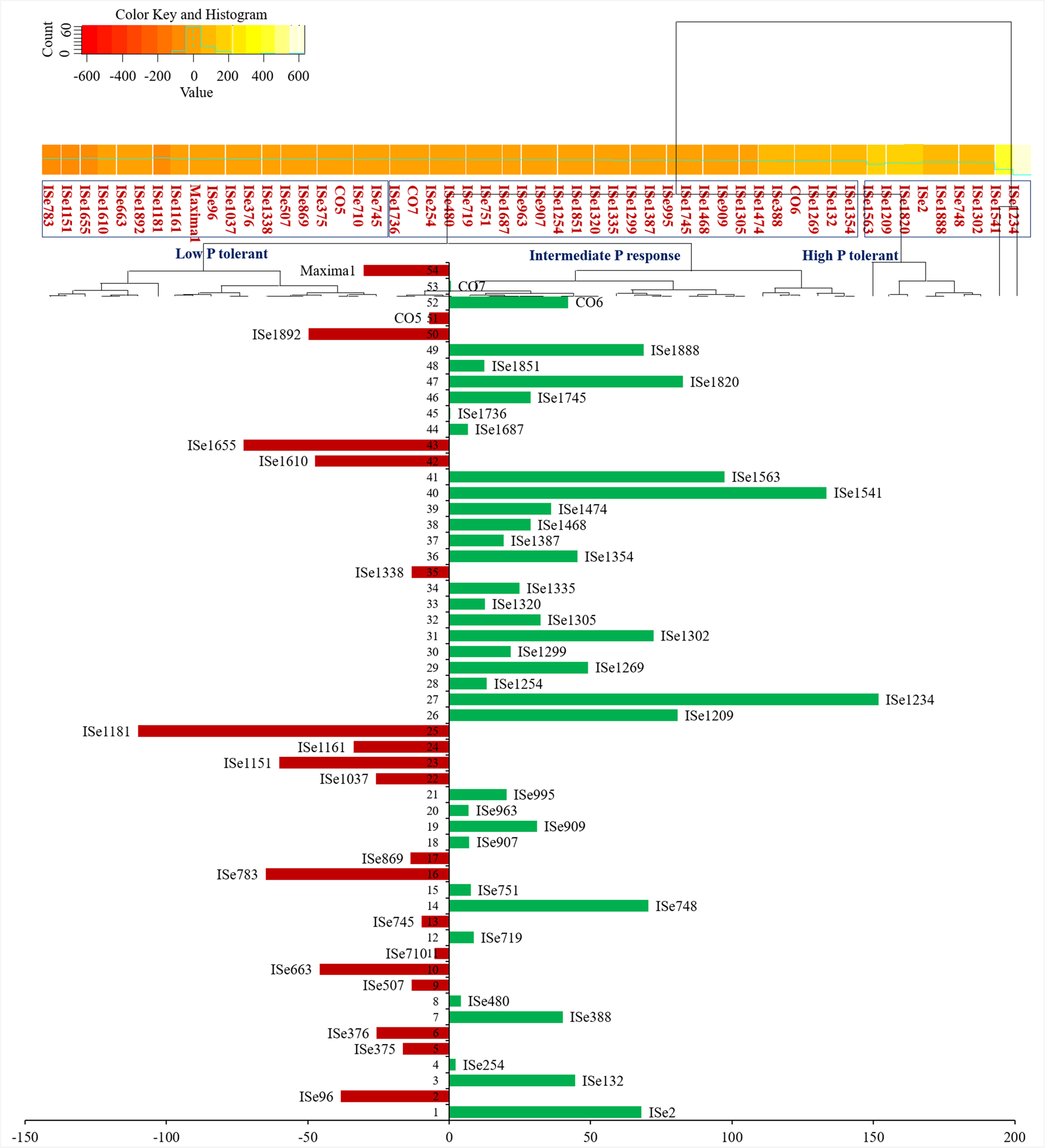
Differential response of genotypes for total seed yield. The genotypes on the left side of the figure (Scale 0 to −150 and red colour bars) tend more towards low P tolerance. Genotypes with long bars on right side of the figure (Scale 0 to 200 and green colour bars) are P+ responders, those closer to the axis are more plastic to P level variations and show stable performance under both P+ and P-conditions.

### Genotype clusters based on P level response in field and greenhouse experiments

The standardised phenotypic data-based analysis of 54 foxtail millet genotypes in both greenhouse and field conditions under P-and P+, revealed two to four clusters of genotypes by *k*-means clustering. The cluster pattern was displayed as a heatmap in **Supplementary Figures S1, S2, S3 and S4**, where yellow equates to a high value and red to a low one. Comparing performance in the field, some of these varieties showed that variabilities in agronomical traits patterned the grouping of foxtail millet genotypes in response to Pi conditions. There were four clusters of genotypes under field conditions in P-level (**Supplementary Figure S1)**. The genotypes ISe 1181, ISe 375 and ISe 376 were low P responders for total seed yield, and these three genotypes were found grouped together into one cluster (**Supplementary Figure S1**). But, these three genotypes were grouped in separate clusters based on the bootstrap probability (BP) and the approximately unbiased (AU) *p*-value (**Figure 7**). The P+ responders ISe 1387, ISe 1687, ISe 663, ISe 869 and CO7, were clustered with genotypes which showed an intermediate response to the unfertilized soil (**Supplementary Figure S1)**. The yield-related traits such as NPT, TSE and SPC showed higher variabilities while the remaining traits such as PH, LF, LL, NC and NL exhibited lesser variabilities in the analysis (**Supplementary Figure S1 and S2)**. We observed similar variations in the analysis for genotypes grown in P-fertilized soil. The most notable example was the genotype ISe 1387 on the top line of the supplementary figure S2, and it was placed in a separate node. This genotype performed relatively well under both greenhouse and field conditions in both phosphate conditions (**Figure 1A, 5** and **Supplementary Figure S2**). The low performers Maxima1, ISe 1037 and ISe 1335 were grouped at the bottom of figure 7 with genotypes ISe 1541 and ISe 1234 which were P+ responders for total seed yield (**Figure 7**).

**Figure 7.**
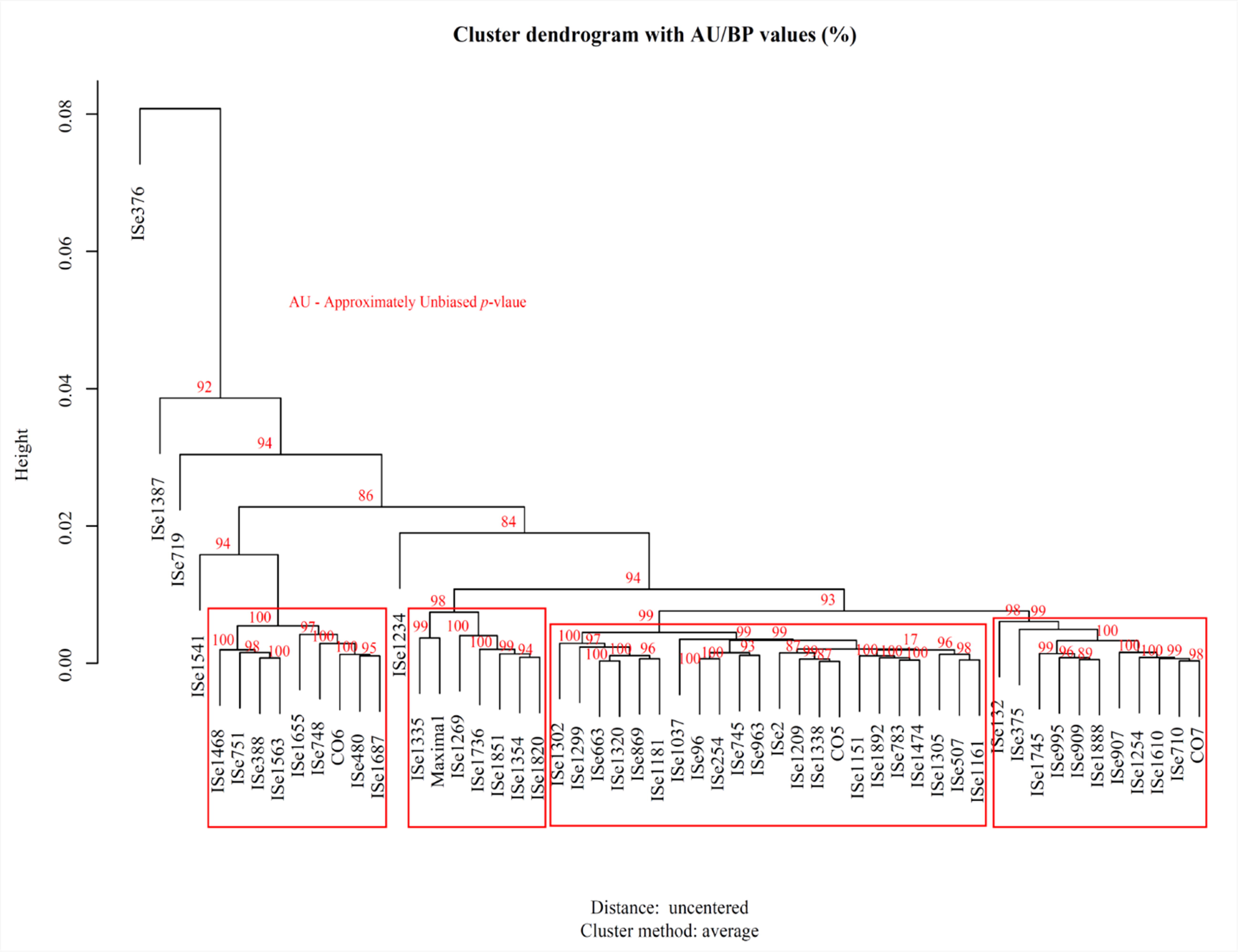
Hierarchical cluster showing the approximately unbiased probability (AU) for the P response genotypes. The value greater than 95% indicates that clusters crated by the variables of P response and node is strongly supported by data. Red boxes indicate the clustering of the genotypes.

The traits measured under greenhouse conditions under P+ treatment (**Supplementary Figure S3)** showed that higher variability in biomass influenced the genotypic clustering pattern more than the remaining traits SL, RL, RHD and RHL that showed lesser variability. Under P-conditions (**Supplementary Figure S4)**, however, the biomass and SL traits were the major traits that diversified genotypes than RHD and RHL. Three groups in the foxtail millet genotypes were identified under P+ condition, with the genotypes CO5, CO6, ISe 2 and Maxima forming a single cluster and are recognised as high performers under P+ (**Supplementary Figure S3).** Most of the low performers were grouped with genotypes that were intermediate in response to the Pi conditions. Within this cluster, we could see all of the high performers (CO5, CO6, ISe 2 and Maxima1) under P+ which gave us confidence in the analysis. Whereas, the traits measured under P- (**Supplementary Figure S4)** created only two groups in the foxtail millet genotypes, and SL showed higher variability. The RHD and RHL showed the same variability in both P- and P+ with good responders, such as ISe 1851, CO6, ISe 1338, ISe 1305 and ISe 254, which were found grouped with genotypes which showed above medium response under P- (**Figure 1B** and **Supplementary Figure S4)**. The genotype Maxima1 was clustered in a separate node and these genotypes showed an intermediate response in P-.

## Discussion

Evaluation of the P response of foxtail millet genotypes under field conditions is important because little information on intrinsic P response of foxtail millet is available and there is potential for translation in practical agriculture. Field situations are complicated by environmental and edaphic factors and biotic influences of the co-existing flora and fauna, however root system responses and rhizosphere environment are difficult to observe and measure. Conversely, greenhouse studies allow evaluation under relatively controlled conditions, and with closer observations, but what plants experience under natural environments such as light, extreme ranges of temperature, nutrient availability and biotic interactions are not recreated under greenhouse environment. Greenhouse based measurements of root system parameters, plant biomass by dry weight and shoot length were used to complement field data to have a comprehensive evaluation of the genotype performance.

The agromorphologic data obtained showed high levels of variation for P response as expected in the field trials. Although many genotypes gave consistent response under both the seasons, a few showed variations due to environmental variations. This can well be attributed to genotypic homeostasis that varies between genotype to genotype. Moreover, this is not unexpected with the current panel of genotypes that are obtained from different countries of origin (Upadhyaya *et al*., 2008). This effect was more apparent with the line Maxima1 which belongs to the race Maxima which prevails in eastern China, Georgia (Eurasia), Japan and Korea. Since the variation imposed was largely due to P input, the remaining effects either through genotypic or through genotype x environment interaction were assumed to be uniform in both the experimental systems employed in this study. Therefore, the genotypes were ranked for growth in terms of higher values of different traits measured. The rank sums were then used as the criteria for selection of extreme genotypes to divulge the contrast in response to P nutrition in foxtail millet genotypes. The ranking is a simple but efficient non-parametrical method in genotype selection (Kang, 1988; Huehn, 1996) but lacks statistical properties. Therefore, to ensure statistical significance between selected individuals, only those traits which showed statistical significance at 95% confidence level were used for ranking, a rank sum calculated and extreme 5% genotypes were selected based on the rank sum. Since the traits of measurement differed under both field and greenhouse screens, the rank sums were computed separately. This identified extreme genotypes under two systems and under two levels of P nutrition. Some of the genotypes such as ISe 1687, Maxima1 and ISe 1387 showed contrasting ranking patterns under field and greenhouse evaluation, which are to be considered as more unstable genotypes, wherein the pattern of P response in one system cannot be construed for the response under the other. Using this context, however, no stable genotype could be identified that was ranked at extreme under both field and greenhouse conditions. The exception to this was ISe 1851, which performed well under both P+ and P-conditions in the field and P-condition in the greenhouse.

P within the plant is translocated easily between the source and sink. The genotypes that possess an active sink of P are P use efficient. There was good agreement between plant P content under field and greenhouse conditions indicating that the P response pattern of the genotypes was consistent irrespective of the growing systems. However, the relationship did not hold for extreme genotypic groups, because of the high magnitude of specific P responses in these sets of genotypes. P uptake efficient genotypes can efficiently harness P under P+ conditions. Furthermore, good responders under P+ and P-conditions accumulated more P in the plant system than the poor responders. Those with high P in the plant system also have better utilisation efficiency if they can effectively trade P between different utilization points of growth and metabolism. Hence, in this experiment good responders under P-can be construed as low P tolerant genotypes, while good responders under P+ can be identified as P loving genotypes. Low P tolerant genotypes with good seed yield such as ISe 663 and ISe 869 are highly preferred for future breeding programmes in foxtail millet targeting P starvation tolerance. Further, the extensive genetic variation for P responses observed in this study, signal the possibility of genetic improvement of foxtail millet for P use efficiency as seen in other cereal crops such as rice (Wissuwa & Ae, 2001), wheat (Batten, 1986), barley (Gahoonia & Nielsen, 2004) and maize (Zhu *et al*., 2005).

Under Pi insufficiency conditions plants reallocate resource to roots increasing the root-shoot ratio and increasing RHD and RHL to promote top soil foraging for P in the growth substrates because more P is available in this layer due to the presence moisture and organic matter, increased microbial activity and oxygen availability (Lynch & Brown, 2001; Péret *et al*., 2014). In soil, root proliferation in response to low P (Péret *et al*., 2014) has been reported in rice conditioned by the QTL, *Pup1* (Wissuwa *et al*., 2005; Heuer *et al*., 2017). Root traits are considered important targets for breeding for better nutrient acquisition (Gahoonia & Nielsen, 2004; Wissuwa, 2003). The formation of root hairs has been considered one of the key strategies to overcome Pi starvation response with minimal carbon cost (Lynch & Ho, 2005). In *Arabidopsis*, increase in number and length of root hairs improved the Pi uptake under Pi-limiting conditions (Bates and Lynch, 2000). All the high responding foxtail millet genotypes (ISe 1851, CO6, ISe 1338, ISe 1305 and ISe 254) produced abundant long root hairs under P- in the greenhouse experiment. RHL and RHD were associated with higher Pi content under P-conditions and Pi content is a good indicator to gain the better information on plant response under low P conditions.

Higher biomass in the genotypes which produced more and longer root hairs is an indication of better P utilization. However, having a high number of root hairs per root was not sufficient. Genotype ISe 1387 had good root hairs under P-conditions but produced extremely small biomass in the glasshouse experiment. However it was the best performer in the field experiment. Other factors such as efficient symbioses or P acquisition ability from organic sources may influence its potential biomass and effect efficient uptake of Pi from the soil. Some foxtail genotypes had very little or minute root hairs produced under P+ conditions, which suggested that proliferation of root hairs was a low P response. In barley, a root hairless mutant reduced the phosphate uptake under the P-condition and was also associated with decreased biomass production (Gahoonia & Nielsen, 2003; Gahoonia *et al*., 2001). The good responders under P-condition in the study had both high RHD and RHL. However, in some crops such as soybean, RHD and RHL under low P conditions showed a negative association (Wang *et al*., 2004). This could be considered as an adaptive response in foxtail millet, a cereal with the adventitious root system, as against the tap root system of soybean wherein a trade-off in terms of carbon use efficiency has been maintained, because combining both RHL and RHD will be too costly in terms of carbon usage. However, in common bean, both higher RHL and RHD were found in P-efficient genotypes (Yan *et al*., 2004).

In the field-based study, the high responding genotypes under unfertilized conditions did almost as well as the top performers under P-fertilized conditions, suggesting high phosphate acquisition and/or use efficiency of these genotypes. Studies of pearl millet and sorghum showed enormous genetic variation for PUtE in natural field conditions based on their responses under P starvation (Leiser *et al*., 2014; Gemenet *et al*., 2014). The changes in the growth, biomass and yield have been recognized as important indicators of Pi deficiency tolerance as reported in other cereals including oat (Andersson *et al*., 2003), rice (He *et al*., 2003; Wissuwa *et al*., 2005), maize (Mollier & Pellerin, 1999) and sorghum (Yoneyama *et al*., 2007).

Since the seed yield is the ultimate target of any crop based study, plasticity analysis of P response was limited to this trait. Plasticity is the difference between P+ and P-response of any genotype under a given cultural condition expressed as the percentage over the average of that genotype in both the conditions. Since the response under low P is subtracted from high P, positive plasticity indicates better performance under fertilised conditions and vice-versa. The genotypes ISe 1387 and ISe 1687 performed well in the natural field in P-fertilized and unfertilized conditions. However, ISe 1387 and ISe 1687 are P+ responders for TSE based on plasticity analysis, because they were more productive under fertilised condition. However, the plasticity of these genotypes is low and closer to the axis (19.3% and 6.6% respectively), which indicated that they could perform equally well under low as well as high P conditions, irrespective of the magnitude of yield, with a slight edge towards high P response. Such genotypes are highly preferable, as they can be grown both under varying P situations.

Clustering analysis and representation of the data as heat maps allowed an overview of the performance of all genotypes under both greenhouse and field conditions. Performances in the greenhouse and in the field were not well correlated in the clusters, and no convincing correlation between any greenhouse measured parameter and seed yield in unfertilized conditions could be ascertained. However, seed yield under P-fertilized and unfertilized conditions was correlated, which suggested that genetic control of seed yield was largely independent of P supply. The total and Pi contents of high-responding genotypes were much higher than that of low responding genotypes, under both the P-fertilized and unfertilized field.. Although all the genotypes grown in the natural field were able to grow in the under unfertilized soil, their uptake efficiency varied considerably resulting in growth differences. This was reflected in contrast between good and poor responding genotypes.

This data is a pertinent reminder that conditions in the field are very different from those in more controlled greenhouse conditions and therefore such data should be used as complementary to the field data to draw conclusions that are practically useful. In an earlier study on rice genotypes grown under natural soil and solution trials, none of the parameters was found to be significantly correlated between soil and solution experiments for relative P-use efficiency (Ni *et al*., 1998). Further, the importance of field-based testing in two different seasons is emphasised to obtain an unbiased performance of genotypes. Several factors like pH of the soil, microorganisms, presence of cations like calcium and aluminium, organic matter substances and mycorrhizal colonisation can influence the availability of Pi to the plants (Gemenet *et al*., 2016). Genotypes that produce fewer root hairs might be compensated for by mycorrhizal colonisation in the field. A recent meta-analysis of mycorrhizal colonisation and responsiveness across a range of crop plants and wild relatives concluded that both colonisation and responsiveness vary markedly (Lehmann *et al*., 2012). While the investigation of interaction with mycorrhizal fungi was beyond the scope of this study it is an important consideration for future study, as is the investigation of root traits such as root branching and angle which are important traits for nutrient efficient root systems (York & Lynch, 2015; Zhu *et al*., 2005).

## Abbreviations

P: Phosphorous
Pi: Inorganic phosphate
PUE: P use efficiency
PAE: Pi acquisition efficiency
QTL: Quantitative trait loci
PUtE: Pi utilization efficiency
RCBD: Randomized complete block design
SDW: Shoot dry weight
RDW: Root dry weight
SL: Shoot length
RL: Root length
RHD: Root hair density
RHL: Root hair length
PH: Plant height
NT: Number of tillers
NPT: Number of productive tillers
NL: Leaf number
LL: Leaf length
LF: Flower length
NC: Cluster number
SPC: Seeds per cluster
25SW: 25 seed weight
TSE: Total seed yield
REML: Restricted maximum likelihood

## Details of supplementary data

**Supplementary Table S1.** Details of foxtail millet genotypes used in the present study

**Supplementary Table S2.** The raw data, the difference in mean values and plasticity analysis between Pi fertilized (P+) and unfertilized (P-) treatments of seedlings grown in natural field condition.

**Supplementary Table S3.** The difference in mean values between P+ (300 µM Pi) and P-(10 µM Pi) treatments of seedlings grown in greenhouse condition

**Supplementary Figure S1.** A clustered heatmap for 54 genotypes of foxtail millet response under unfertilized (P-) soil. Genotypes are scaled and hierarchically clustered by Euclidean distance. Yellow equates to a high value and red to a low value. PH, plant height; NPT, productive tiller number; NL, leaf number; LL, leaf length; LF, flower length; NC, cluster number; SPC, seeds per cluster; TSE total seed yield.

**Supplementary Figure S2.** A clustered heatmap for 54 genotypes of foxtail millet response under Pi fertilized (P+) soil. Genotypes are scaled and hierarchically clustered by Euclidean distance. Yellow equates to a high value and red to a low value. PH, plant height; NPT, productive tiller number; NL, leaf number; LL, leaf length; LF, flower length; NC, cluster number; SPC, seeds per cluster; TSE total seed yield.

**Supplementary Figure S3.** A clustered heatmap for 54 genotypes of foxtail millet response for the greenhouse under P+ (300 µM Pi). Genotypes are scaled and hierarchically clustered by Euclidean distance. Yellow equates to a high value and red to a low value. SL, shoot length; RL, root length; RHD, root hair density; RHL, root hair length.

**Supplementary Figure S4.** A clustered heatmap for 54 genotypes of foxtail millet response for the greenhouse under P-(10 µM Pi). Genotypes are scaled and hierarchically clustered by Euclidean distance. Yellow equates to a high value and red to a low value. SL, shoot length; RL, root length; RHD, root hair density; RHL, root hair length.

### Acknowledgements

We thank the Tamil Nadu Agricultural University (TNAU), Coimbatore, Tamil Nadu, India, for supplying the seeds of three local genotypes of foxtail millet. We thank TNAU at Regional Research Station (RRS), Paiyur, Tamil Nadu, India, and the field workers Mr Palani and his family members, Ms Vadamalli, Ms Kalaivani, Mr, Murugan, and others at Paiyur, Tamil Nadu, India for the helps with the field study.

## Conflict of interest statement

The authors declare that there is no conflict of interest.

## Authors and Contributors

Conceived and designed the experiments: SAC, MR, SI, AB; Performed the experiments: SAC, MR, VR; Analyzed the data: SAC, MR, KKV; Contributed reagents/materials/analysis tools: HDU, SI, AB; Wrote the paper: SAC, MR, KKV, AB, SI, HDU.

