## Supplementary figures and images for "Diverse phenotypic responses and phosphate content in foxtail millet genotypes under greenhouse and field conditions"

### Supplementary Figure S1

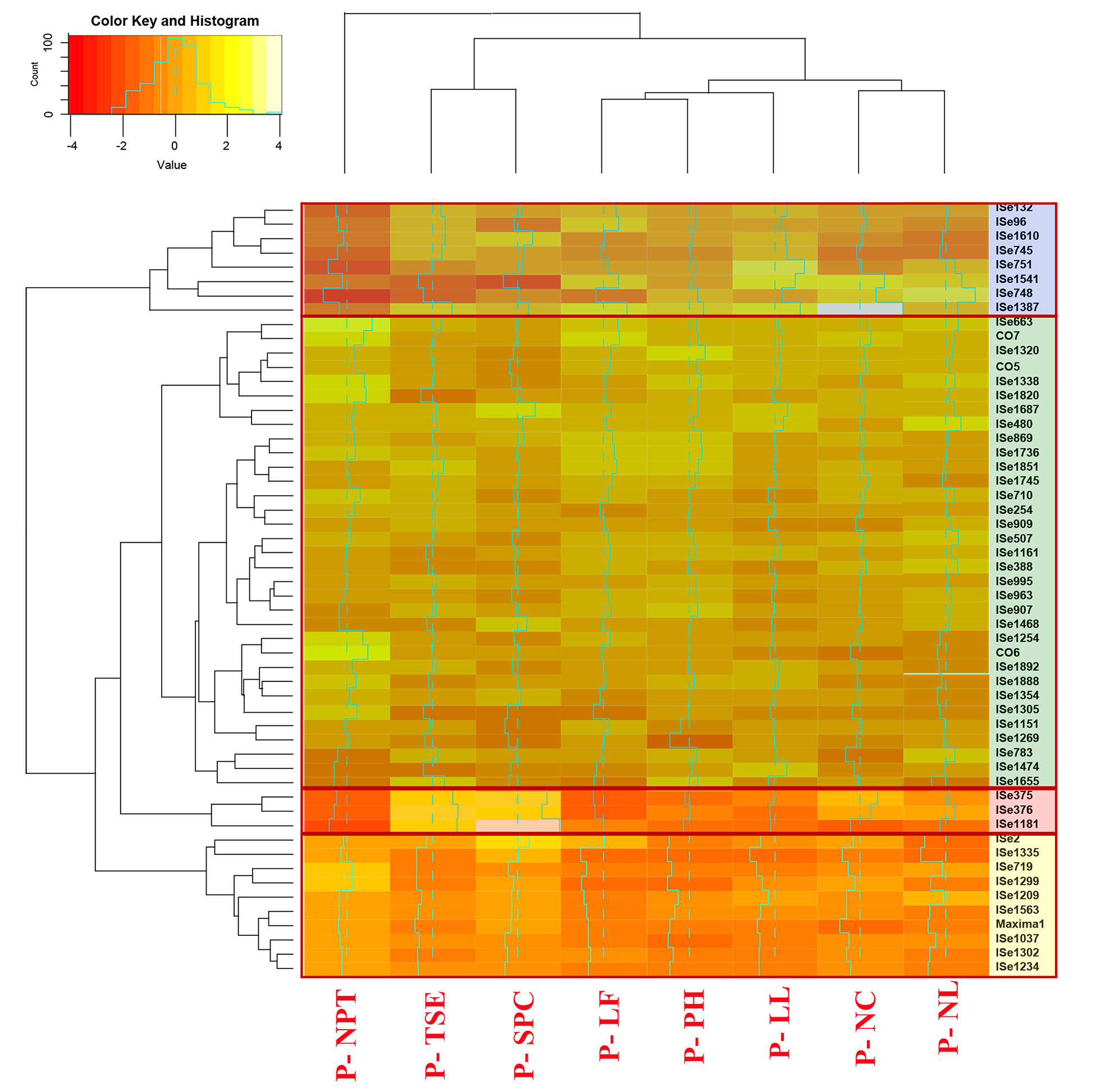

### Supplementary Figure S2

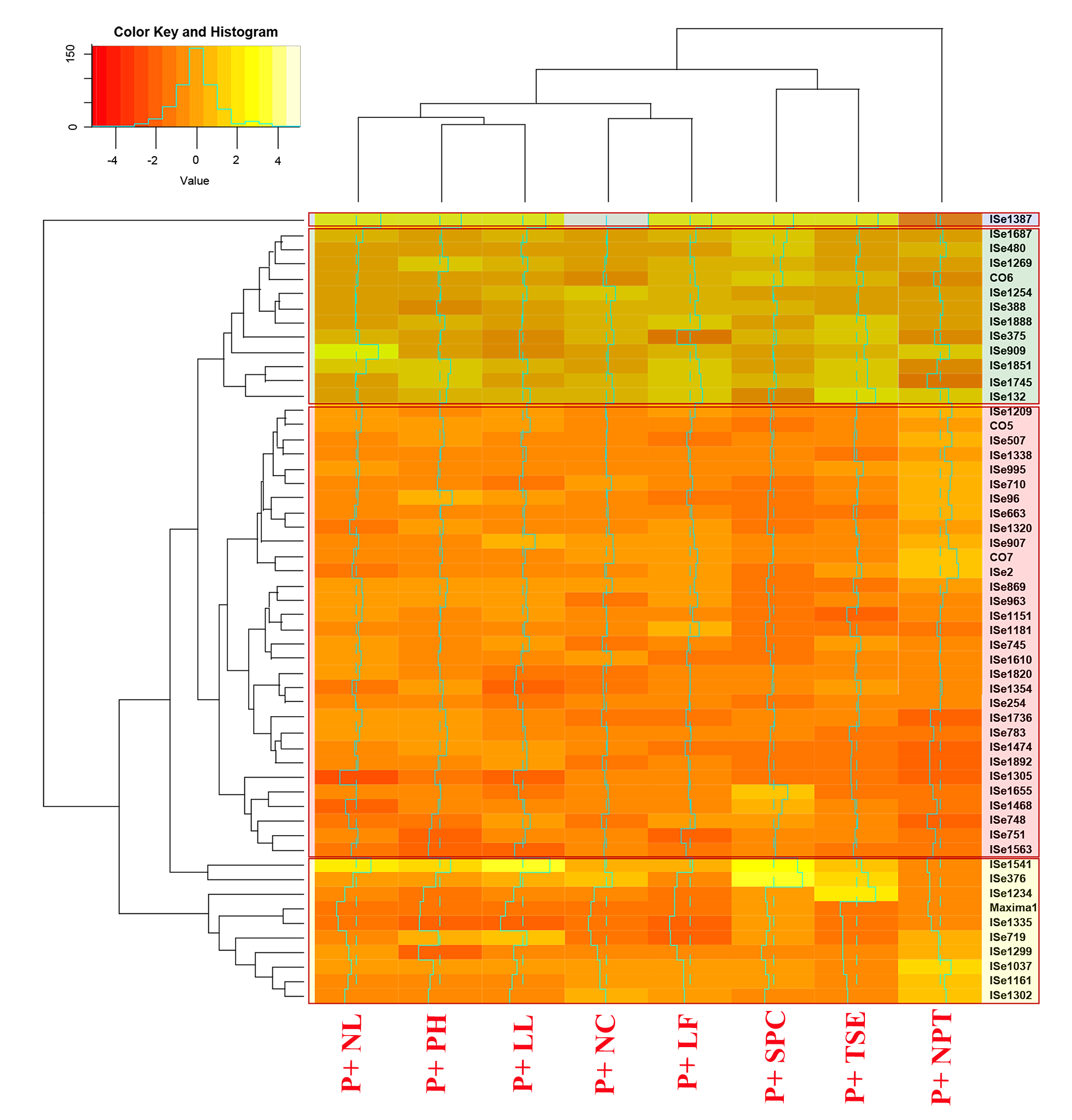

### Supplementary Figure S3

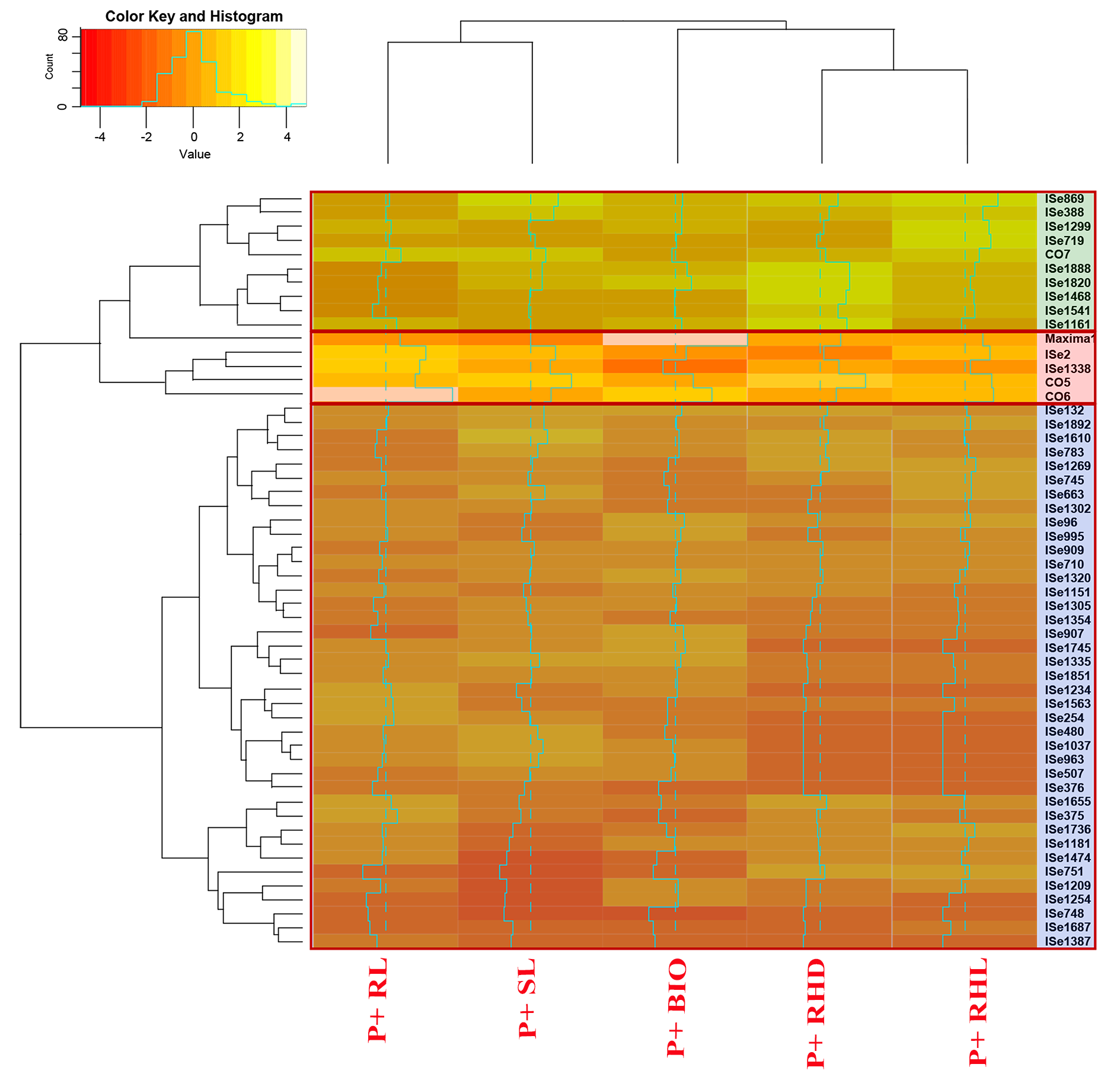

### Supplementary Figure S4

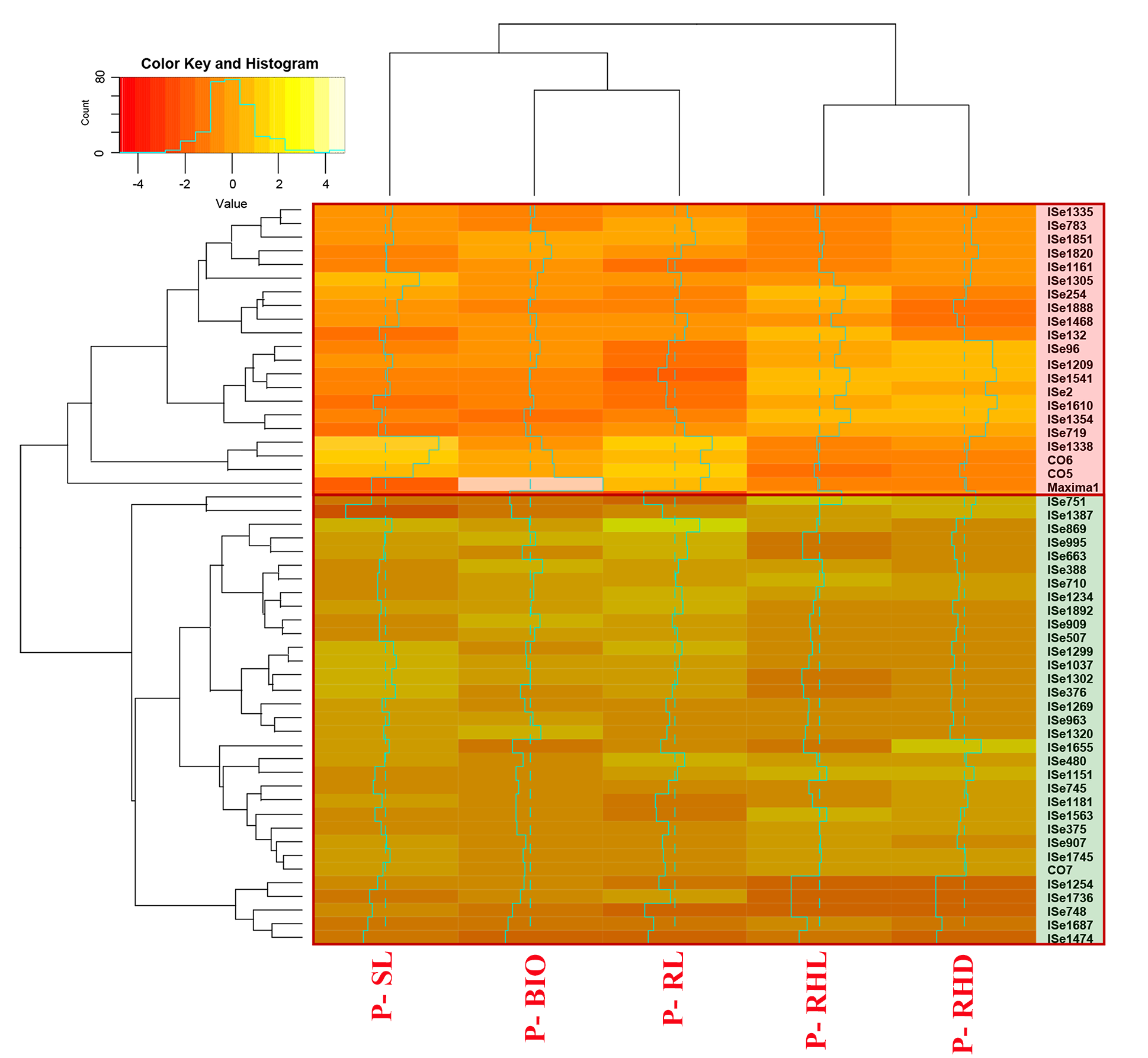
