## Supplementary tables S1 and S2 for "Diverse phenotypic responses and phosphate content in foxtail millet genotypes under greenhouse and field conditions"

**Supplementary Table S1.** Details of foxtail millet genotypes used in the present study

| **S. No** | **Name of the genotype** | **Alternate Name/ Number** | **Origin** |
| --- | --- | --- | --- |
| 1 | ISe 2 | Korra | India |
| 2 | ISe 96 | Bhedi | India |
| 3 | ISe 132 | Kangani | India |
| 4 | ISe 254 | Mobbu navne | India |
| 5 | ISe 375 | Koni dhan | India |
| 6 | ISe 376 | - | India |
| 7 | ISe 388 | Tangum | India |
| 8 | ISe 480 | Var.hungsheku | China |
| 9 | ISe 507 | M 235; B 35 | Kenya |
| 10 | ISe 663 | FAO ≠ 11324 | Switzerland |
| 11 | ISe 710 | - | India |
| 12 | ISe 719 | - | Pakistan |
| 13 | ISe 745 | - | India |
| 14 | ISe 748 | - | India |
| 15 | ISe 751 | - | India |
| 16 | ISe 783 | - | India |
| 17 | ISe 869 | Rgot | India |
| 18 | ISe 907 | SE 201 | India |
| 19 | ISe 909 | SE 480 | India |
| 20 | ISe 963 | SE 3045 | India |
| 21 | ISe 995 | SE 7230/3-1 | India |
| 22 | ISe 1037 | IPM 1626 | Lebanon |
| 23 | ISe 1151 | NESE 82: 3876-1 | Syria |
| 24 | ISe 1161 | NESE 87-2: 3881-2 | Syrian Arab Republic |
| 25 | ISe 1181 | EC 130490 | China |
| 26 | ISe 1209 | EC 131210: WIR 864 | Russia & CISs |
| 27 | ISe 1234 | EC 131236: WIR 1030 | Russia & CISs |
| 28 | ISe 1254 | EC 131256: WIR 1346 | Russia & CISs |
| 29 | ISe 1269 | EC 134287 | South Africa |
| 30 | ISe 1299 | EC 134283; PI 250025 | Iran |
| 31 | ISe 1302 | EC 134286; Pi 207502 | Afghanistan |
| 32 | ISe 1305 | PI 283988; EC 134289 | Spain |
| 33 | ISe 1320 | EC 134329; PI 363068 | USA |
| 34 | ISe 1335 | EC 134320; PI 290461 | Hungary |
| 35 | ISe 1338 | EC 134257; PI 173805 | Turkey |
| 36 | ISe 1354 | - | India |
| 37 | ISe 1387 | EC 135535 | Sri Lanka |
| 38 | ISe 1468 | SIA 326 | India |
| 39 | ISe 1474 | EC 155192-1 | United kingdom |
| 40 | ISe 1541 | T 132/12 | India |
| 41 | ISe 1563 | Wooljin 7 | Republic of Korea |
| 42 | ISe 1610 | - | Malawi |
| 43 | ISe 1655 | 40051 | Taiwan |
| 44 | ISe 1687 | Jhum | India |
| 45 | ISe 1736 | Acc No. 6542 | Nepal |
| 46 | ISe 1745 | Shwe sutkon | Myanmar |
| 47 | ISe 1820 | ISe 231 A; Wari | India |
| 48 | ISe 1851 | ISe 275 A: Kangani | India |
| 49 | ISe 1888 | ISe 410 B; Roll boer mann | Ethiopia |
| 50 | ISe 1892 | ISe 472 A | USA |
| 51 | CO-5 | - | India |
| 52 | CO-6 | - | India |
| 53 | CO-7 | - | India |
| 54 | Maxima | Bs 3875 | - |

**Supplementary Table S2.** The difference in mean values between P+ (300 µM Pi) and P- (10 µM Pi) treatments of seedlings grown in greenhouse condition

| **S. No** | **Name of the genotype** | **Biomass** | **SL** | **RL** | **RHD** | **RHL** |
| --- | --- | --- | --- | --- | --- | --- |
| 1 | ISe 2 | 2.03 | 8.80 | 8.67 | -58.67 | -0.51 |
| 2 | ISe 96 | 0.50 | 0.87 | 1.43 | -74.33 | -0.54 |
| 3 | ISe 132 | 0.63 | 6.90 | -1.50 | -25.00 | -0.65 |
| 4 | ISe 254 | -1.20 | -0.50 | 0.47 | -44.00 | -0.80 |
| 5 | ISe 375 | -0.03 | 0.37 | 3.33 | -25.33 | -0.31 |
| 6 | ISe 376 | -0.47 | -0.90 | -1.67 | -24.00 | -0.19 |
| 7 | ISe 388 | -0.17 | 9.70 | -0.67 | -13.67 | -0.20 |
| 8 | ISe 480 | 0.03 | 4.53 | -2.37 | -44.67 | -0.38 |
| 9 | ISe 507 | -0.67 | 2.97 | -1.77 | -29.67 | -0.28 |
| 10 | ISe 663 | 0.70 | 6.60 | -3.53 | -22.00 | 0.02 |
| 11 | ISe 710 | 0.03 | 3.80 | -0.73 | -23.33 | -0.33 |
| 12 | ISe 719 | 1.17 | 4.60 | -1.60 | -64.33 | -0.40 |
| 13 | ISe 745 | 0.40 | 3.40 | 1.37 | -32.33 | -0.08 |
| 14 | ISe 748 | -0.77 | -3.30 | 3.50 | 0.00 | 0.00 |
| 15 | ISe 751 | 0.13 | -4.40 | 2.80 | -43.67 | -0.58 |
| 16 | ISe 783 | 0.83 | 3.33 | -5.40 | -37.33 | -0.25 |
| 17 | ISe 869 | 1.60 | 9.20 | -4.57 | -6.67 | -0.02 |
| 18 | ISe 907 | 2.17 | 2.50 | 0.17 | -29.00 | -0.35 |
| 19 | ISe 909 | -0.50 | 4.30 | -0.33 | -20.33 | -0.11 |
| 20 | ISe 963 | 0.07 | 4.03 | 0.63 | -26.33 | -0.24 |
| 21 | ISe 995 | 0.57 | -0.13 | -2.23 | -20.67 | -0.05 |
| 22 | ISe 1037 | 0.63 | 4.30 | -1.23 | -27.00 | -0.25 |
| 23 | ISe 1151 | 2.00 | 1.83 | -0.93 | -47.00 | -0.45 |
| 24 | ISe 1161 | -0.37 | 2.13 | 3.47 | -22.33 | -0.28 |
| 25 | ISe 1181 | 2.10 | -3.33 | 3.80 | -37.00 | -0.15 |
| 26 | ISe 1209 | 0.00 | -5.63 | 0.90 | -81.67 | -0.52 |
| 27 | ISe 1234 | 0.30 | -0.43 | -0.50 | -34.33 | -0.37 |
| 28 | ISe 1254 | 1.67 | -3.73 | 0.13 | 1.67 | 0.04 |
| 29 | ISe 1269 | 0.27 | 3.47 | 1.20 | -4.67 | -0.04 |
| 30 | ISe 1299 | 1.97 | 0.60 | -0.50 | -10.33 | -0.01 |
| 31 | ISe 1302 | -0.50 | 1.57 | -0.50 | -19.00 | 0.01 |
| 32 | ISe 1305 | -0.20 | -4.10 | -3.43 | -44.67 | -0.53 |
| 33 | ISe 1320 | -0.10 | 2.17 | 0.57 | -7.67 | -0.09 |
| 34 | ISe 1335 | 0.70 | 3.67 | -2.17 | -57.33 | -0.28 |
| 35 | ISe 1338 | -2.66 | -0.27 | -2.10 | -25.67 | -0.20 |
| 36 | ISe 1354 | 0.63 | 2.13 | -1.77 | -72.67 | -0.78 |
| 37 | ISe 1387 | 0.23 | 2.73 | 1.20 | -51.67 | -0.42 |
| 38 | ISe 1468 | -0.37 | -0.03 | -3.83 | -3.00 | -0.38 |
| 39 | ISe 1474 | 1.13 | -1.17 | 5.03 | 9.13 | 0.01 |
| 40 | ISe 1541 | 0.47 | 1.50 | 1.33 | -65.00 | -0.66 |
| 41 | ISe 1563 | 1.23 | 1.37 | 5.30 | -35.33 | -0.45 |
| 42 | ISe 1610 | 0.57 | 8.93 | 0.53 | -72.67 | -0.51 |
| 43 | ISe 1655 | 0.93 | -1.37 | 3.93 | -51.00 | -0.04 |
| 44 | ISe 1687 | 0.40 | 0.00 | 1.90 | -8.30 | -0.19 |
| 45 | ISe 1736 | 0.67 | -0.26 | 0.43 | 9.33 | 0.21 |
| 46 | ISe 1745 | 2.80 | 1.57 | 2.53 | -42.67 | -0.44 |
| 47 | ISe 1820 | -0.13 | 5.33 | -2.63 | -31.33 | -0.29 |
| 48 | ISe 1851 | -1.07 | 1.27 | -4.70 | -48.67 | -0.41 |
| 49 | ISe 1888 | 2.30 | 3.77 | -0.80 | 4.67 | -0.55 |
| 50 | ISe 1892 | 0.50 | 6.63 | -1.43 | -11.00 | -0.14 |
| 51 | CO5 | -0.20 | 9.73 | -2.50 | 7.00 | -0.02 |
| 52 | CO6 | 2.47 | 1.46 | 5.90 | -23.67 | -0.08 |
| 53 | CO7 | 0.70 | 7.07 | 4.60 | -30.00 | -0.18 |
| 54 | Maxima | 1.00 | 4.50 | -3.10 | -20.00 | -0.13 |

^SL, shoot length; RL, root length; RHD, root hair density, RHL, root hair length^
